# SARS-CoV-2 RNA shedding in recovered COVID-19 cases and the presence of antibodies against SARS-CoV-2 in recovered COVID-19 cases and close contacts

**DOI:** 10.1101/2020.07.17.208439

**Authors:** Chintana Chirathaworn, Manit Sripramote, Piti Chalongviriyalert, Supunee Jirajariyavej, Phatharaporn Kiatpanabhikul, Jatuporn Saiyarin, Chulikorn Soudon, Orawan Thienfaidee, Thitisan Palakawong Na Ayuthaya, Chantapat Brukesawan, Dootchai Chaiwanichsiri, Duangnapa Intharasongkroh, Nasamon Wanlapakorn, Jira Chansaenroj, Jiratchaya Puenpa, Ritthideach Yorsaeng, Arunee Thitithanyanont, Rungrueng Kitphati, Anek Mungaomklang, Pijaya Nagavajara, Yong Poovorawan

**Author notes:** Department of Microbiology, Faculty of Medicine, Chulalongkorn University, Bangkok 10330 Thailand (Chintana Chirathaworn). Division of Academic Affairs, Faculty of Medicine, Chulalongkorn University (Nasamon Wanlapakorn). These authors contributed equally to this work (Chintana Chirathaworn, Manit Sripramote). Share last authorship. E-mail address (YP).

## Abstract

Coronavirus disease 2019 (COVID-19) is caused by severe acute respiratory syndrome coronavirus 2 (SARS-CoV-2). COVID-19 emerged in December 2019 and has spread globally. Although Thailand has been effective at controlling the spread of COVID-19, disease surveillance and information on antibody responses in infected cases and close contacts are needed because there is still no specific treatment or vaccine available. We investigated 217 recovered COVID-19 cases to monitor their viral RNA shedding and production of antibodies against SARS-CoV-2. The presence of antibodies in blood samples from 308 close contacts of COVID-19 cases was also determined. Viral RNA was still detectable in 6.6 % of recovered COVID-19 cases. The most prolonged duration of viral RNA shedding detected in this study was 105 days. IgM, IgG, and IgA antibodies against SARS-CoV-2 were detected in 13.82, 88.48, and 83.41 % of the recovered cases 4–12 weeks after disease onset, respectively. Although the patients had recovered from their illness, the levels of antibodies detected showed association with their symptoms during their stay in hospital. Fifteen of the 308 contacts (4.87 %) of COVID-19 cases tested positive for IgG antibodies. The presence of antibodies against SARS-CoV-2 suggested that there was viral exposure among close contacts. Viral clearance and the pattern of antibody responses in infected individuals are both crucial for effectively combatting SARS-CoV-2. Our study provides additional information on the natural history of this newly emerging disease related to both natural host defenses and a strategy for vaccine development.

## Introduction

Coronavirus disease 2019 (COVID-19) is an emerging infectious disease caused by a novel coronavirus named severe acute respiratory syndrome coronavirus 2 (SARS-CoV-2). The disease emerged in December 2019 and has since spread globally. On March 11, 2020, the World Health Organization (WHO) declared the coronavirus outbreak to be a pandemic. Currently, over 11 million cases of COVID-19 have been reported worldwide, resulting in more than 500,000 deaths [1]. The number of confirmed COVID-19 cases reported in Thailand was 3,195, with 58 deaths [2]. In Thailand, the first case of COVID-19 was detected in the capital, Bangkok, in mid-January 2020. During the first few months, most of the reported cases in Thailand were associated with travelers who had visited other countries. However, the number of reported cases increased rapidly owing to the spreading of the disease *via* entertainment venues and Thai boxing stadiums.

SARS-CoV-2 is an enveloped, positive-sense single-stranded RNA virus of the Coronaviridae family. It was classified as a novel betacoronavirus similar to the previously identified SARS-CoV (severe acute respiratory syndrome coronavirus) and MERS-CoV (Middle East severe respiratory syndrome coronavirus) [3,4]. SARS-CoV-2 enters host cells through the binding of its spike glycoprotein to the host angiotensin-converting enzyme 2 (ACE2) receptor. This virus can spread across species. The reproductive number (R_0_) of SARS-CoV-2 is higher than that of SARS-CoV and MERS-CoV, suggesting that this novel coronavirus has a greater pandemic potential [5–7]. Although the number of reported COVID-19 cases in Thailand has decreased rapidly [2], the surveillance of infected cases and disease transmission is crucial for disease prevention before a vaccine can be developed.

SARS-CoV-2-infected patients present with a wide range of symptoms, ranging from mild to severe. Severe illness includes pneumonia, difficulty breathing, and respiratory failure. SARS-CoV-2-infected individuals can also be asymptomatic [8].

Real-time reverse transcription-polymerase chain reaction (RT-PCR) for viral RNA detection is widely used for the laboratory-based diagnosis of COVID-19. Antibody detection has also been increasingly reported to indicate exposure to SARS-CoV-2. Although several reports have suggested the combined use of antibody and viral RNA detection, real-time RT-PCR is currently the method-of-choice for SARS-CoV-2 diagnosis [9–11].

Viral RNA has been shown to be detectable in recovered patients, although the duration of viral RNA shedding reported varied among infected patients. Viral RNA shedding can last at least 6 weeks [12–16]. Liu et al. reported that patients with severe symptoms had a higher viral load and a longer viral shedding period than patients with mild symptoms (<10 days for the group with mild symptoms and >10 days for the group with severe symptoms). These authors investigated viral RNA in nasopharyngeal (NP) swabs and the delta Ct values were used to represent the viral load [17].

Antibody responses following infection can be utilized for disease diagnosis and can also represent evidence of viral exposure. Detection of IgM, IgG, and IgA antibodies against SARS-CoV-2 antigens in COVID-19 have been reported. Although antibody detection is currently not widely used for COVID-19 diagnosis, the presence of antibodies provides additional valuable information. The presence of antibodies is indicative of immune responses to SARS-CoV-2, immune clearance, and pathogenesis in infected patients. Additionally, the presence of antibodies reveals the transmission of the virus to contacts.

In this study, we followed recovered COVID-19 cases using real-time RT-PCR to investigate the duration of viral shedding as detected in NP swabs. Antibody detection was performed to monitor antibody levels in all the recovered patients. Blood samples from individuals who had a history of contact with COVID-19 patients were also collected for antibody detection.

## Materials and methods

The study protocol was approved by the Institutional Ethics Committee (IRB) of the Bangkok Metropolitan Administration, Thailand (IRB No. M001h/63_Exp). The IRB waived the need for consent because the used samples were obtained from routine preventive measures and were de-identified and anonymous.

### Samples and sample collection

Samples included in this study were collected from recovered COVID-19 patients and their close contacts. The recovered COVID-19 cases recruited in this study were patients who had been diagnosed with SARS-CoV-2 infection and admitted to hospitals or public health centers under the Bangkok Metropolitan Administration. NP swabs and blood samples were collected for the detection of viral RNA and determination of the presence of antibodies, respectively. Close contacts were individuals with a history of contact with COVID-19 patients. Only blood samples for antibody detection were collected from close contacts.

For recovered cases, the duration between the first day of symptom onset to the day of sample collection was recorded. For close contacts, the duration between the last contact and sample collection was noted.

### Real-time RT-PCR for the detection of viral RNA

Real-time RT-PCR targeting the RNA-dependent RNA polymerase (RdRp), envelope (E), and nucleocapsid (N) genes of SARS-CoV-2 was performed using the LightMix Modular SARS and Wuhan CoV E-gene Kit (TIB MOBIOL, Synheselabor GmbH, Berlin, Germany) in a LightCycler 480 II system following the manufacturer’s recommendation (Roche Diagnostics International Ltd., Rotkreuz, Switzerland). The results were reported as detectable or undetectable based on Ct values.

### Detection of antibodies against SARS-CoV-2 by enzyme-linked immunoassays

For recovered COVID-19 cases, IgM, IgG, and IgA antibodies against SARS-CoV-2 were determined, whereas, for close contacts, IgG antibodies were investigated in blood samples. IgM antibody levels were measured using the MAGLUMI 2000 fully automatic chemiluminescent analytical system (Snibe, Shenzhen, China) according to the manufacturer’s instructions. The values were reported as arbitrary unit per milliliter (AU/mL). The determination of IgG and IgA antibodies was performed using an ELISA automated system (Euroimmun, Luebeck, Germany) following the manufacturer’s instructions. The optical density (OD) at 450 nm was measured and the ratio was calculated using the reading of each sample to the reading of the calibrator. The result was obtained as a ratio between sample and cut-off OD value. Sample to cut off value of > 1 was considered seropositive.

### Statistical analysis

The association between antibody levels and clinical characteristics was analyzed using the Mann–Whitney *U*-test and chi-square test. The difference between the number of people positive for IgG antibodies against SARS-CoV-2 in each group of close contacts was determined using the chi-square test. A *p*-value <0.05 was considered statistically significant.

## Results

A total of 217 recovered COVID-19 cases were recruited in this study. Based on the patients’ records during their stay in hospital, the clinical manifestations of the recovered COVID-19 cases were considered to be asymptomatic (4/217), mild (151/217), presenting with pneumonia (59/217), and presenting with pneumonia with tracheal intubation (3/217). A total of 308 close contacts were included in this study. Of these, 118 were household contacts, while the rest were close friends, colleagues, health care personnel who took care of the COVID-19 cases, taxi drivers, neighbors, or individuals who lived or performed activities in the same community as the COVID-19 cases.

All samples were collected from April to June 2020. Gender, age, and other information relating to the recovered COVID-19 cases and close contacts gathered in this study are shown in Table 1.

**Table 1.** Recovered COVID-19 cases and close contacts included in this study and the results of antibodies detection.

|  | Recovered cases | Close contacts |
| --- | --- | --- |
| **Total number** | 217 | 308 |
| **Age (years)** | 2–76 | 2–77 |
| **Gender** |  |  |
| Number of males (age in years) | 92 (2–76) | 159 (2–69) |
| Number of females (age in years) | 125 (11–70) | 149 (7–77) |
| **History of symptoms** |  |  |
| Mild/asymptomatic; number (%) | 155/217 (71.43) | NA |
| Pneumonia; number (%) | 59/217 (27.19) |  |
| Pneumonia with intubation; number (%) | 3/217 (1.38) |  |
| **Duration from symptom onset to sample collection (days)** | 28–142 | NA |
| **Duration from the last contact to sample collection (days)** | NA | 1–128 |
| **IgM antibody detection** |  |  |
| Number of positive cases (%) | 30/217 (13.82) | ND |
| Duration from symptom onset to sample collection for positive cases (days) | 34–67 |  |
| **IgG antibody detection** |  |  |
| Number of positive cases (%) | 192/217 (88.48) | 15/308 (4.87) |
| Duration from symptom onset to sample collection for positive case (days) | 28–142 |  |
| **IgA antibody detection** |  |  |
| Number of positive cases (%) | 181/217 (83.41) | ND |
| Duration from symptom onset to sample collection for positive cases (days) | 28–142 |  |

NA: not applicable; ND: not done.

### Detection of SARS-CoV-2 RNA in recovered COVID-19 cases

NP swabs could not be obtained for 5 of the 217 recovered COVID-19 cases recruited in this study. Among the 212 cases whose NP swabs were collected, 14 (6.6 %) showed detectable viral RNA by real-time RT-PCR. The ages of these 14 cases ranged from 16–67 years. The Ct values obtained from the real-time RT-PCR analysis indicated the low amount of viral RNA left. RT-PCR analysis of NP swabs from 2 of the 14 recovered cases showed Ct values of 28.43 and 29.61. The Ct values obtained from the NP swabs of the other 12 cases ranged between 30.22 and 37.74. The duration between the day of first onset and sample collection ranged from 36– 105 days (mean = 57 days). Our study suggested that viral RNA shedding was detectable for up to 15 weeks.

### Antibodies against SARS-CoV-2 in recovered COVID-19 cases

The levels of IgM, IgG, and IgA antibodies against SARS-CoV-2 in blood samples from 217 recovered cases were measured. Fig 1A–C depicts the analysis of the data from the determination of the three antibody isotypes against SARS-CoV-2 according to the duration (in weeks) of the first day of symptom onset to the day of sample collection. Among the 217 cases, 30 (13.82 %), 192 (88.48 %), and 181 (83.41 %) were positive for IgM, IgG, and IgA antibodies, respectively. All 30 IgM-positive samples were also positive for IgG and IgA antibodies. In addition, 150/217 (69.12 %) cases were positive for both IgG and IgA antibodies. The duration between the day of the first symptom onset and blood sample collection varied from 28–142 days. As shown in Table 1, although only 13.82 % of recovered cases were IgM-positive, IgM antibodies remained detectable for up to 2 months in some cases. IgG and IgA antibodies were detectable for up to 20 weeks.

**Fig 1.**
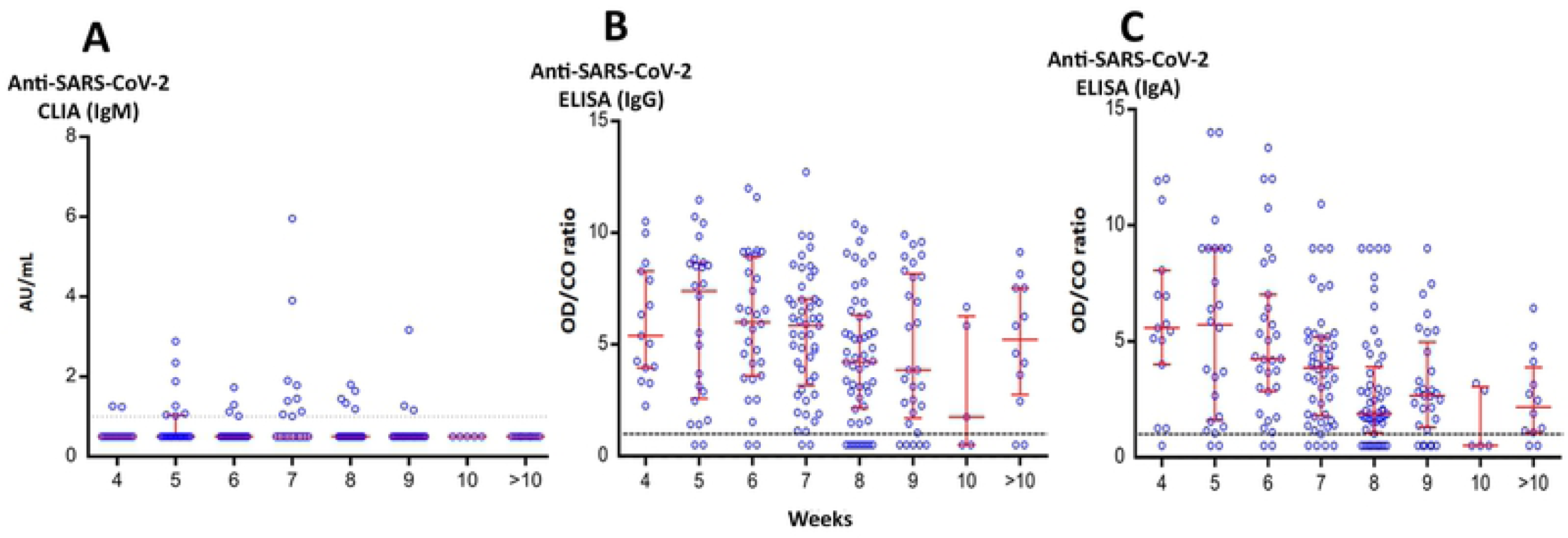
Antibodies against SARS-CoV-2 in recovered COVID-19 cases. IgM (A), IgG (B), and IgA (C) antibodies against SARS-CoV-2 in recovered COVID-19 cases were determined as mentioned in the Materials and methods. The data represent the results according to the duration (weeks) between the first day of symptom onset and the day of sample collection. Bars represent median values (middle line) and the upper and lower interquartile range (IQR) (upper and lower lines).

As mentioned above, COVID-19 symptoms included asymptomatic, mild, pneumonia, and pneumonia with tracheal intubation. To investigate the association of patient symptoms with the findings of this study, patient symptoms were grouped into nonpneumonia (asymptomatic and mild symptoms; 155/217 cases, mean age = 34.5) and pneumonia (62/217 cases, mean age = 42.2) groups. The ages of cases were significantly different between the groups with and without pneumonia (chi-square, *p* = 0.027). Moreover, when the association between antibody concentrations and symptoms was analyzed, the levels of IgM, IgG, and IgA antibodies against SARS-CoV-2 were significantly higher in the pneumonia group than in the nonpneumonia group (*p* = 0.001, *p* < 0.0001, and *p* < 0.0001, respectively) (Fig 2A–C).

**Fig 2.**
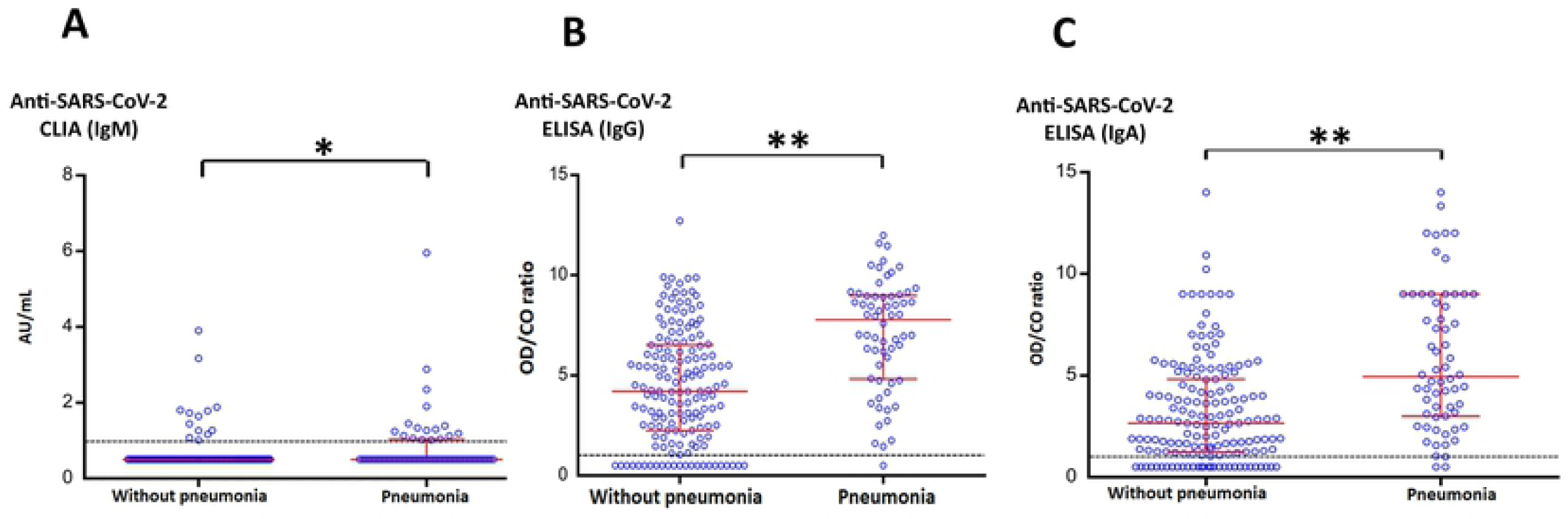
The association between the antibodies against SARS-CoV-2 and symptoms of recovered COVID-19 cases. The levels of IgM (A), IgG (B), and IgA (C) antibodies against SARS-CoV-2 in recovered COVID-19 cases with and without pneumonia are shown. Bars represent median values (middle line) and 1× the upper and lower interquartile range (IQR) (upper and lower lines). The levels of IgM, IgG, and IgA antibodies against SARS-CoV-2 were higher in patients with pneumonia than in those without pneumonia. * *p* = 0.0002, ** *p* < 0.00001.

### Antibody detection in close contacts of SARS-CoV-2 patients

IgG antibodies against SARS-CoV-2 detected in close contacts of COVID-19 patients are shown in Fig 3A. Among the 308 close contacts, 15 (4.87 %) were positive for IgG antibodies, and two of these 15 close contacts presented with respiratory symptoms. Blood samples from healthy donors collected in 2018, i.e., before the emergence of COVID-19, were tested for the presence of antibodies against SARS-CoV-2. All 50 samples showed negative results.

**Fig 3.**
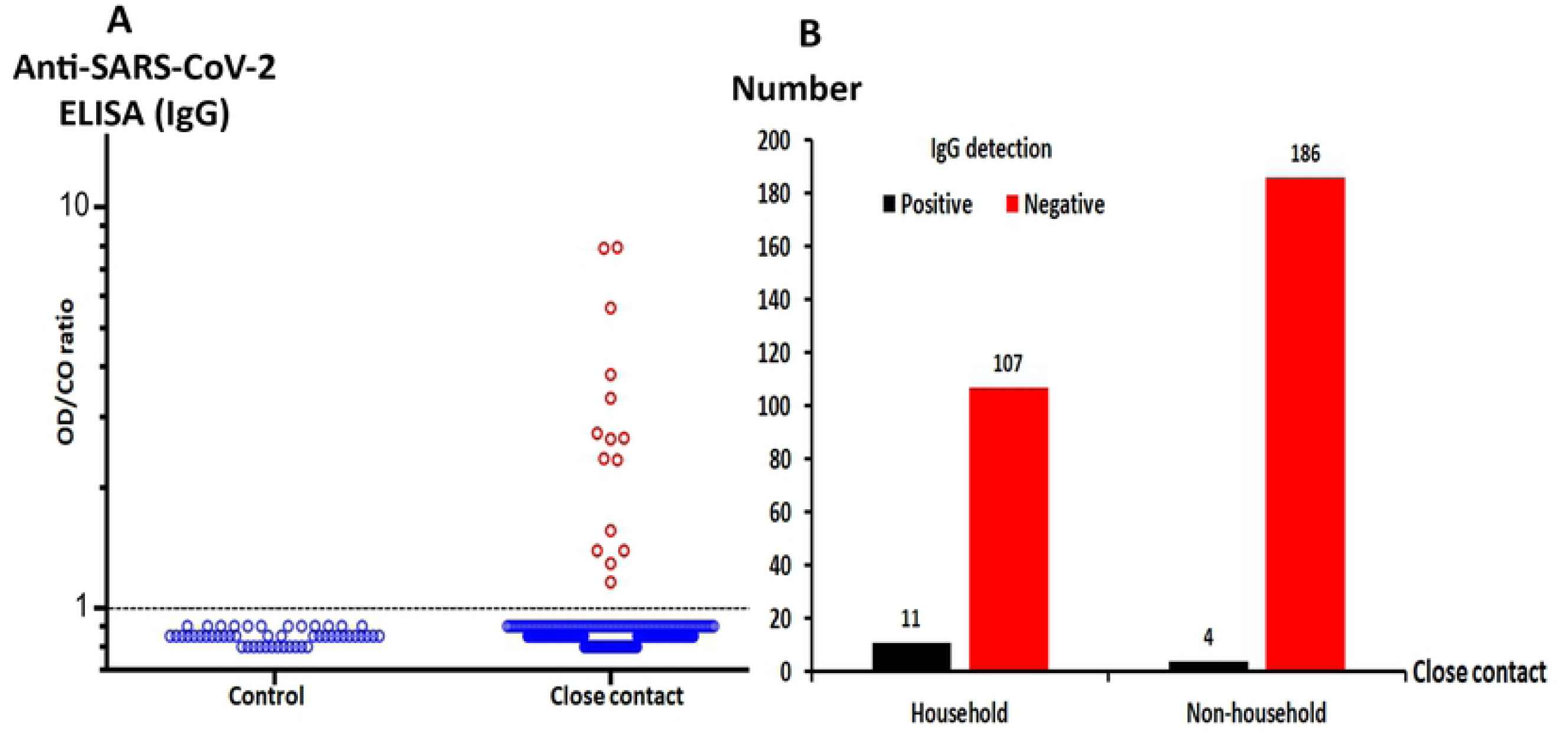
IgG antibodies against SARS-CoV-2 in close contacts of COVID-19 cases. IgG antibodies against SARS-CoV-2 in blood samples from 308 close contacts of COVID-19 cases and 50 healthy controls collected in 2018 were determined. IgG antibodies were detected in 15/308 close contacts, but none of the healthy controls (A). The numbers of IgG-positive household and nonhousehold contacts are shown (B).

Of the 15 IgG-positive contacts, 11 (73.33 %) were household contacts and 4 (26.67 %) were colleagues and/or friends of the COVID-19 cases (Fig 3B). Our data supported that household contacts represent a high-risk group for disease exposure. A total of 118 household contacts were recruited in this study. Our results showed that the prevalence of viral exposure among the household contacts, as evidenced by the presence of antibodies, was 9.3 % (11 out of 118 contacts). The prevalence of viral exposure among the other types of close contacts was 2.1 % or 4 out of 190 contacts (chi-square, *p* = 0.015).

## Discussion

The number of cases of COVID-19 has increased rapidly worldwide since the emergence of SARS-CoV-2 infection in December 2019. Laboratory diagnosis and treatments have been urgently evaluated. In addition, the duration of viral RNA shedding and patterns of immune responses, especially antibody production, have been studied. The report of a COVID-19 case in Taiwan showed that SARS-CoV-2 RNA could be detected on day 5 after disease onset. IgM antibodies have also been detected on day 11, and are still detectable on day 27 [18]. Xiang et al. demonstrated the dynamics of antibody responses in samples collected from COVID-19 cases at 3–40 days after symptom onset. The authors reported that IgM and IgG antibodies against SARS-CoV-2 could be detected as early as the fourth day after symptom onset. In confirmed cases, the sensitivity of the IgM and IgG tests were 77.3 and 83.3 %, respectively [19]. Nisreen et al. assessed the levels of IgG and IgA antibodies in one severe and two mild cases of COVID-19, and found that the concentrations of these antibodies were higher in severe cases. Samples were collected between 6 and 27 days after diagnosis, and the samples from the severe case showed earlier and higher seroconversion [20]. The same finding was demonstrated in MERS-CoV infection [21,22]. A study on SARS-associated coronavirus showed that IgG antibodies persist longer than IgA antibodies and are therefore more suitable for disease surveillance [23]. A different report from Finland showed that the neutralizing antibodies, IgM and IgG, could be detected within 9 days of symptom onset. The antibodies were still detectable on day 20 after symptom onset [24].

Long et. al. reported that IgG and IgM antibody concentrations were higher in severe cases of COVID-19, while IgM levels decreased slightly after the third week of symptom onset. This study followed up 63 patients by collecting serum at 3-day intervals, and 96.8 % of the patients showed seroconversion. In addition, 26 patients who were initially negative for IgG and IgM antibodies were followed up for the detection of seroconversion. The median number of days for IgG and IgM seroconversion was 13 days after symptom onset. Both IgG and IgM antibodies were detected in the in 9/26 samples, IgM was detected earlier than IgG in 7/26 patients, and later in 10/26 cases [25]. Yongchen et al. reported the longitudinal follow-up by viral RNA and antibody testing of 5 severe, 11 non-severe, and 3 asymptomatic COVID-19 cases. Viral RNA remained detectable for 9–33 days (median = 14), 2–21 days (median = 10), and 5–28 days (median = 5) in severe, non-severe, and asymptomatic cases, respectively. All symptomatic patients in this study showed antibody responses. Three patients showed seroconversion within the first week. Moreover, this study showed that the antibody to SARS-CoV-2 was detectable for at least 6 weeks. However, only 20 % (1/5) of asymptomatic cases showed an antibody response, with this one patient showing seroconversion in the third week after diagnosis [26]. Zhao et al. reported the dynamics of total, IgM, and IgG antibodies against SARS-CoV-2 with disease progression in 173 COVID-19 cases. Less than 40 % of the patients showed detectable antibodies within the first week after symptom onset. On day 15 after disease onset, 100 %, 94.3 %, and 79.8 % of cases were positive for total, IgM, and IgG antibodies, respectively. IgM appeared before IgG (median time to seroconversion was 12 and 14 days, respectively). This study showed that the sensitivity of IgM antibody detection was higher than for IgG antibodies. Viral RNA detection showed the greatest sensitivity during the first week of symptom onset. In addition, this study showed that high antibody titers were associated with the worst clinical manifestations [27].

Close contacts of COVID-19 cases have been previously investigated. Bi et al. investigated 391 COVID-19 cases and 1,286 close contacts, and showed that household contacts and individuals who traveled with COVID-19 cases had a high risk of infection [28]. Meanwhile, Guo et al. showed that the median duration of antibody detection was 5 days for IgM and IgG and 14 days for IgG. In this study, samples were collected between 1–39 days after disease onset. These authors also investigated four close contacts of a family with 2 COVID-19 cases, and found that these 4 close contacts were asymptomatic. IgM antibodies were detected in 3 of the 4 close contacts, whereas IgG antibodies were undetectable [29].

In the current study, we demonstrated that viral shedding could still be detected in 14/212 (6.6 %) of the recovered COVID-19 cases. The longest duration of viral shedding shown in our study was 105 days. Several reports have indicated that the antibody levels persisted for at least 6 weeks. We chose to follow the recovered cases to investigate the persistence of antibodies against SARS-CoV-2. The IgM level, which commonly declines rapidly following infection, was still detectable in 13.82 % of recovered cases. IgA, an immune component of mucosal immunity, and IgG were still detectable in more than 80 % of recovered cases. It would be interesting to investigate further the recovered cases whose antibodies against SARS-CoV-2 were undetectable.

We also found that high levels of antibodies were associated with disease severity. Interestingly, although we investigated the antibody levels in the recovered cases, an association between high antibody levels and severe cases was still observed. This suggests that patients with high antibody levels could have a higher viral burden. However, there is still no clear evidence to indicates that a correlation exists between the number of antibodies produced and viral load. A high number of antibodies should facilitate viral clearance and alleviate disease symptoms. However, the roles of the antibodies produced in COVID-19 cases remain to be elucidated.

In this study, we investigated a large group of close contacts, and found that 4.8 % of them displayed evidence of viral exposure. Besides, our results showed that household contact represent a high-risk group for disease exposure which is essential for the prevention of disease transmission.

In conclusion, our study provided information on COVID-19 cases after disease recovery. The data suggested that further monitoring should be performed to achieve a full understanding of the duration of viral clearance and antibody responses. Our study, which investigated both recovered cases and close contacts, was intended to support the policy followed by Thailand on the prevention of the spread of SARS-CoV-2 infection in the country. Our findings regarding the high antibody response in recovered patients will be useful for the recruitment of volunteers by the National Blood Center for the donation of convalescent plasma. Additionally, our data are useful for the further understanding of COVID-19 transmission and infection control, while our results on the patterns of antibody production and duration of antibody detection may inform strategies for achieving herd immunity and as well as for vaccine immunization.

## Acknowledgments

We are grateful to the staff of the Center of Excellence in Clinical Virology for their technical and administrative assistance. We greatly appreciate the recovered Covid-19 cases and their close contacts in Thailand for their kind contribution and collaboration. With all their helps, the interesting information obtained from this study could be gathered for future development of Covid-19 therapeutic and vaccine strategies.

## References

1. WHO. Coronavirus disease 2019 (COVID-19). Situation Report-167. Available at hwwidd-scs-r-c--s-ps. Accessed July 6 (2020).

2. Department of Disease Control MoPH, Thailand. Covid-19 Situation Reports. Available at, https://covid19.ddc.moph.go.th/en Accessed July 6 (2020).

3. Tu YF, Chien CS, Yarmishyn AA, Lin YY, Luo YH, Lin YT, et al. A Review of SARS-CoV-2 and the Ongoing Clinical Trials. Int J Mol Sci. 2020;21(7). doi: 10.3390/ijms21072657. PMID: 32290293.

4. Coronaviridae Study Group of the International Committee on Taxonomy of V. The species Severe acute respiratory syndrome-related coronavirus: classifying 2019-nCoV and naming it SARS-CoV-2. Nat Microbiol. 2020;5(4):536–44. doi: 10.1038/s41564-020-0695-z. PMID: 32123347

5. Petrosillo N, Viceconte G, Ergonul O, Ippolito G, Petersen E. COVID-19, SARS and MERS: are they closely related? Clin Microbiol Infect. 2020;26(6):729–34. doi: 10.1002/14651858.CD013582. PMID: 32315451

6. Shang J, Wan Y, Luo C, Ye G, Geng Q, Auerbach A, et al. Cell entry mechanisms of SARS-CoV-2. Proc Natl Acad Sci U S A. 2020;117(21):11727–34. doi: 10.1073/pnas.2003138117. PMID: 32376634

7. Wu JT, Leung K, Leung GM. Nowcasting and forecasting the potential domestic and international spread of the 2019-nCoV outbreak originating in Wuhan, China: a modelling study. Lancet. 2020;395(10225):689–97. doi: 10.1016/S0140-6736(20)30260-9. PMID: 32014114

8. Lauer SA, Grantz KH, Bi Q, Jones FK, Zheng Q, Meredith HR, et al. The Incubation Period of Coronavirus Disease 2019 (COVID-19) From Publicly Reported Confirmed Cases: Estimation and Application. Ann Intern Med. 2020;172(9):577–82. doi: 10.7326/M20-0504. PMID: 32150748

9. Liu W, Liu L, Kou G, Zheng Y, Ding Y, Ni W, et al. Evaluation of Nucleocapsid and Spike Protein-based ELISAs for detecting antibodies against SARS-CoV-2. J Clin Microbiol. 2020 May 26;58(6):e00461–20. doi: 10.1128/JCM.00461-20. PMID: 32229605

10. Pan Y, Li X, Yang G, Fan J, Tang Y, Zhao J, et al. Serological immunochromatographic approach in diagnosis with SARS-CoV-2 infected COVID-19 patients. J Infect. 2020 Jul;81(1):e28–e32. doi: 10.1016/j.jinf.2020.03.051. PMID: 32283141

11. Li Z, Yi Y, Luo X, Xiong N, Liu Y, Li S, et al. Development and clinical application of a rapid IgM-IgG combined antibody test for SARS-CoV-2 infection diagnosis. J Med Virol. 2020. doi: 10.1002/jmv.25727. PMID: 32104917

12. Qi L, Yang Y, Jiang D, Tu C, Wan L, Chen X, et al. Factors associated with duration of viral shedding in adults with COVID-19 outside of Wuhan, China: A retrospective cohort study. Int J Infect Dis. 2020. doi: 10.1016/j.ijid.2020.05.045. PMID: 32425636

13. Fu Y, Han P, Zhu R, Bai T, Yi J, Zhao X, et al. Risk Factors for Viral RNA Shedding in COVID-19 Patients. Eur Respir J. 2020 Jul 2;56(1):2001190. doi: 10.1183/13993003.01190-2020. PMID: 32398298.

14. Qian GQ, Chen XQ, Lv DF, Ma AHY, Wang LP, Yang NB, et al. Duration of SARS-CoV-2 viral shedding during COVID-19 infection. Infect Dis (Lond). 2020;52(7):511–2. doi: 10.1080/23744235.2020.1748705. PMID: 32275181.

15. Zhou F, Yu T, Du R, Fan G, Liu Y, Liu Z, et al. Clinical course and risk factors for mortality of adult inpatients with COVID-19 in Wuhan, China: a retrospective cohort study. Lancet. 2020;395(10229):1054–62. doi: 10.1016/S0140-6736(20)30566-3. PMID: 32171076.

16. Liu WD, Chang SY, Wang JT, Tsai MJ, Hung CC, Hsu CL, et al. Prolonged virus shedding even after seroconversion in a patient with COVID-19. J Infect. 2020 Apr 10;S0163-4453(20)30190-0. doi: 10.1016/j.jinf.2020.03.063. PMID: 32283147.

17. Liu Y, Yan LM, Wan L, Xiang TX, L. A, Liu JM, et al. Viral dynamics in mild and severe cases of COVID-19. Lancet Infect Dis. 2020;20(6):656–7. doi: 10.1016/S1473-3099(20)30232-2. PMID: 32199493.

18. Lee NY, Li CW, Tsai HP, Chen PL, Syue LS, Li MC, et al. A case of COVID-19 and pneumonia returning from Macau in Taiwan: Clinical course and anti-SARS-CoV-2 IgG dynamic. J Microbiol Immunol Infect. 2020 Jun;53(3):485–487. doi: 10.1016/j.jmii.2020.03.003. PMID: 32198005.

19. Xiang F, Wang X, He X, Peng Z, Yang B, Zhang J, et al. Antibody Detection and Dynamic Characteristics in Patients with COVID-19. Clin Infect Dis. 2020 Apr 19. doi: 10.1093/cid/ciaa461. PMID: 32306047.

20. Nisreen MAO, Marcel AM, Wentao L, Chunyan W, Corine HG, Victor MC, et al. Severe Acute Respiratory Syndrome Coronavirus 2−Specific Antibody Responses in Coronavirus Disease 2019 Patients. Emerging Infectious Disease journal. 2020 Jul;26(7):1478–1488. doi: 10.3201/eid2607.200841. PMID: 32267220.

21. Choe PG, Perera R, Park WB, Song KH, Bang JH, Kim ES, et al. MERS-CoV Antibody Responses 1 Year after Symptom Onset, South Korea, 2015. Emerg Infect Dis. 2017;23(7):1079–84. doi: 10.3201/eid2307.170310. PMID: 28585916.

22. Alshukairi AN, Khalid I, Ahmed WA, Dada AM, Bayumi DT, Malic LS, et al. Antibody Response and Disease Severity in Healthcare Worker MERS Survivors. Emerg Infect Dis. 2016;22(6):1113–1115. doi: 10.3201/eid2206.160010. PMID: 27192543.

23. Hsueh PR, Huang LM, Chen PJ, Kao CL, Yang PC. Chronological evolution of IgM, IgA, IgG and neutralisation antibodies after infection with SARS-associated coronavirus. Clin Microbiol Infect. 2004;10(12):1062–6. doi: 10.1111/j.1469-0691.2004.01009.x. PMID: 15606632.

24. Haveri A, Smura T, Kuivanen S, Osterlund P, Hepojoki J, Ikonen N, et al. Serological and molecular findings during SARS-CoV-2 infection: the first case study in Finland, January to February 2020. Euro Surveill. 2020;25(11). doi: 10.2807/1560-7917.ES.2020.25.11.2000266. PMID: 32209163.

25. Long QX, Liu BZ, Deng HJ, Wu GC, Deng K, Chen YK, et al. Antibody responses to SARS-CoV-2 in patients with COVID-19. Nat Med. 2020 Jun;26(6):845–848. doi: 10.1038/s41591-020-0897-1. PMID: 32350462.

26. Yongchen Z, Shen H, Wang X, Shi X, Li Y, Yan J, et al. Different longitudinal patterns of nucleic acid and serology testing results based on disease severity of COVID-19 patients. Emerg Microbes Infect. 2020;9(1):833–6. doi: 10.1080/22221751.2020.1756699. PMID: 32306864.

27. Zhao J, Yuan Q, Wang H, Liu W, Liao X, Su Y, et al. Antibody responses to SARS-CoV-2 in patients of novel coronavirus disease 2019. Clin Infect Dis. 2020 Mar 28. doi: 10.1093/cid/ciaa344. PMID: 32221519.

28. Bi Q, Wu Y, Mei S, Ye C, Zou X, Zhang Z, et al. Epidemiology and transmission of COVID-19 in 391 cases and 1286 of their close contacts in Shenzhen, China: a retrospective cohort study. Lancet Infect Dis. 2020 Apr 27;S1473-3099(20)30287-5. doi: 10.1016/S1473-3099(20)30287-5. PMID: 32353347.

29. Guo L, Ren L, Yang S, Xiao M, Chang, Yang F, et al. Profiling Early Humoral Response to Diagnose Novel Coronavirus Disease (COVID-19). Clin Infect Dis. 2020 Mar 21. doi: 10.1093/cid/ciaa310. PMID: 32198501.

